# Fruit scent chemistry: adaptation to seed dispersal interactions

**DOI:** 10.64898/2026.08.27.744376

**Authors:** Linh M. N. Nguyen, Diary Razafimandimby, Rebekka Sontowski, Darren CJ Wong, Jana Ebersbach, John C. D’Auria, Radoniaina R. Rafaliarison, Kim Valenta, Nicole M. van Dam, Philipp M. Schlüter, Omer Nevo

## Abstract

Fleshy fruits have evolved diverse traits to attract seed dispersers in response to frugivore behavior and sensory capacities. Fruit scent has been suggested to signal ripeness and nutritional quality, yet the volatile components involved and the information they convey remain poorly understood. It is unknown which information is encoded in fruit scent, whether plants actively synthesize these signals, and thus whether scent constitutes an evolved communication system shaping seed-dispersal interactions. Aliphatic esters, chemicals whose odor is often described as “fruity,” are abundant in some ripe fruits, especially those dispersed by animals which tend to rely on their sense of smell for fruit selection. Moreover, they have been argued to be associated with sugar content, potentially rendering them an honest signal and hence a hotspot of animal-plant chemical communication. We investigated whether aliphatic esters indicate fruit quality honestly and represent an adaptive trait shaped by disperser identity. Using 13 fig species (*Ficus* spp.; Moraceae) in Madagascar, we quantified seed dispersal by multiple animals using a quantitative ecological network. We then quantified chemical signals and nutritional rewards using thermal desorption gas chromatography-mass spectrometry (TD-GCMS) and high-performance liquid chromatography (HPLC), and used genome-guided transcriptome assembly to identify the candidate genes responsible for ester signaling. Our results show that (a) aliphatic esters occur more frequently in species dispersed primarily by olfactory-oriented mammals than in those dispersed by visually oriented birds; (b) ester abundance correlates positively with soluble sugar across species only in mammal-dispersed species, indicating an honest signal that is activated only when ecologically relevant; and (c) putative AAT gene revealing elevated expression associated with the increased abundance of aliphatic esters, sugars in single mammal-dispersed taxon. Together, these findings support the hypothesis that fruits have evolved to utilize the biochemical link between esters (signals) and sugars (rewards) to provide honest signals to seed dispersers.

**Significance Statement:** Fruit–frugivore communication is a critical component of ecological systems as it allows animals to find and identify ripe fruit, thus facilitating seed dispersal, plant reproduction, forest regeneration, and animal community maintenance. Chemical communication via fruit scent remains underexplored, particularly regarding the information encoded, active signal synthesis, and adaptive significance. Using a model system of wild figs from Madagascar, our study suggests that fruits have evolved to utilize aliphatic esters, a group of chemicals commonly described as “fruity” by human observers, to signal sugar levels to seed-dispersing animals. In species that rely on lemurs, which tend to rely strongly on their sense of smell, ester levels were higher than in bird-dispersed species, where visual signals are more prominent. Further, in lemur-dispersed species ester levels were correlated with sugar, indicating that they are reliably signaling fruit quality. Linking these patterns to elevated expression of the gene mediating ester biosynthesis, we also identify a molecular mechanism potentially under selection. These findings establish fruit scent as an adaptive communication system and reveal how plant chemicals evolve under mammalian selection.

## Introduction

Seed dispersal is a cornerstone of ecosystem functioning, and animal-mediated dispersal by fruit-eating animals (frugivores) is a predominant strategy among angiosperms, particularly in the tropics (1). Fleshy fruits have evolved repeatedly and independently across most angiosperm families, often in shaded environments that favor larger, energy-rich seeds (2), as such conditions place a premium on attracting effective animal vectors. By consuming fleshy fruits, frugivores obtain nutritional rewards while moving seeds away from the parent plant, helping plants reduce competition and predation near the parent plant and thereby promote recruitment (3).

Fruit traits such as size, color, and nutrient content influence detectability, attractiveness, and handling efficiency, ultimately affecting the probability that a given animal will disperse a plant’s seeds (4–6). Frugivores select fruits in response to this adaptive landscape. In fruit-eating birds, gape width constrains the size of fruits they can swallow: broad-gaped species consume larger fruits on average (7). Fruit bats rely primarily on smell to locate fruit, and bat-dispersed figs can produce scent compounds that attract bats but are rejected by bird-dispersed figs (8). Animals do not remove all fruits equally but preferentially select fruits based on traits, thereby influencing seed dispersal and plant reproductive success. In turn, selection on fruit traits and animal foraging strategies can drive reciprocal diversification and trait matching between fruits and their dispersers, including matching in morphology (9, 10), sensory capacities (11), and foraging behavior (12). This reciprocal relationship underlies the dispersal syndrome hypothesis, which proposes that suites of fruit traits are associated with the morphology, behavior, and sensory capacities of their dispersers. Such interactions may also shape contemporary seed-dispersal networks, although their strength varies among ecological communities (13, 14).

Among these traits, fruit scent, parallel with fruit color (15), has been proposed as a signal for ripeness in plant species that depend on animal dispersers, which tend to rely on their sense of smell for fruit selection (16, 17). Fruit scent, the bouquet of mainly specialized volatiles emitted by the ripe fruit, plays a vital yet often poorly studied role in the interaction between plants bearing fleshy fruits and animal dispersers. Signaling of fruit ripeness has been suggested to manifest in a shift in chemical quantity and diversity in ripe fruits compared to unripe fruits, generating a unique chemical signature marking fruits as ripe (16, 17). In turn, frugivores use fruit scent for multiple purposes, for instance, bats use odor-guided detection together with echolocation (8, 18–23) and primates might use scent signals in pre-ingestive fruit selection by sniffing behavior correlated with fruit olfactory conspicuousness (16, 17, 24, 25).

So far, existing models of scent signaling propose a binary signaling system, where fruit scent indicates only whether a fruit is ripe or unripe (17, 23, 26). Yet fruit scents are composed of a wide array of volatile organic compounds, such as terpenoids, benzenoids, aldehydes and ketones (27). Different classes of volatile compounds are derived from distinct biosynthetic pathways and enzymatic systems, potentially enabling fruits to encode multiple types of information for dispersers while simultaneously fulfilling other ecological or physiological functions (27, 28). However, not all detected scent compounds are necessarily involved in dispersal signaling; some likely serve alternative roles, such as defending against herbivores and pathogens, attracting pollinators (29), and regulating plant development and stress responses (30). This raises the possibility that current approaches introduce substantial noise, including chemically detectable but functionally irrelevant volatiles, which could mask the ecological and evolutionary signals of the compounds that mediate interactions. For these reasons, the specific fruit scent compounds involved in plant–disperser signaling have thus far not been identified.

Several chemicals, specifically volatiles, have been suggested to be associated with nutritional content, making them an honest signal to attract seed dispersers (31, 32). Indeed, volatile profiles are associated with nutritional rewards: while protein levels show no consistent relationship with volatile composition, sugar content strongly predicts fruit scents composition, but only in a few species (33). This link is important for both sides of the interaction: sugars represent the fast energetic reward driving frugivore foraging decisions in frugivores, such as elephants, which can detect sugar-rich fruits by sensing key volatiles such as ethanol and ethyl acetate (25, 31, 34). For plants, however, volatile production requires an investment of carbon that could otherwise be allocated to primary metabolism. This trade-off between cost-benefit asymmetry is the engine of the selection. The signal might not simply be “fruit odor,” but a chemical profile whose information value depends on how reliably it predicts.

Esters are promising candidates for fruit–frugivore signaling because they are volatile at ambient temperatures and often arise from ethanol, which ultimately derives from sugars, making them easily perceivable olfactory cues that could help mammals detect ripe fruits at a distance or assess ripeness at close range (16, 17, 35). Aliphatic esters, strong candidates for genuine dispersal signaling, are often dominant in ripe fruit scents and appear particularly abundant in mammal-dispersed species (17). This raises the possibility that esters provide information about fruit rewards. However, evidence linking esters to sugar content remains limited to a small number of plants–disperser systems. For example, esters have been proposed as cues to sugar rewards in wild tomatoes (36), while studies of other mammal-dispersed fruits have reported associations between aliphatic esters and sugar content (33, 37). Together, the occurrence of aliphatic esters in systems where chemical communication with olfactory-guided dispersers is expected, along with their association with sugar content, suggests that they represent a chemically coherent and ecologically grounded signal within a largely noisy volatile landscape.

Biochemically, components facilitating the ester production include alcohols, acids and alcohol acyltransferases (*AAT*s) enzymes (38, 39). This ester formation reaction depends on the activity of the catalyzed enzyme, and also on the availability of metabolic building blocks (40), which is one of the limiting factors of acyl-CoAs derived from sugar metabolism. Alcohols may originate either from methanol released during cell wall modification due to the action of pectin methylesterases (41) or ethanol generated through sugar fermentation (42, 43); both pathways can ultimately contribute to ester formation, and critically, both are associated with fruit maturation. As such, the substrates for ester formation are directly linked to fruit ripeness and quality. Another component required for ester synthesis is *AAT* expression in the fruit (39, 40, 44, 45). *AAT*s (EC 2.3.1.84) belong to the BAHD acyltransferase family (Pfam: PF02458), with orthologs found across land plants (46). Across multiple species, *AAT* activity correlates consistently with ester production during ripening (47–50). This suggests that variation in scent signals is actively regulated by *AAT* gene expression and substrate availability and hence can be an honest signal. Although evidence for signal-reward linkages in wild fruits remains limited, the available data converge on esters as a focal point for chemical communication (33, 34, 37), making them a compelling candidate for studying how fruit scent mediates plant-frugivore co-evolution.

Aliphatic esters may reflect functions shaped by the selection pressures of costly signaling that drive the evolution of plants. If fruits convert sugar-derived substrates into esters, these compounds could provide honest advertisements that attract dispersers. Frugivores that can detect high-quality fruits gain advantages in intra-group competition by preferring individuals that provide reliable information (37, 51). Aliphatic esters are therefore particularly interesting because they sit at the intersection of three processes: ripening, odor production, and potentially detectable reward chemistry. They are prominent in ripe-fruit scent systems, can be generated through a mechanistical *AAT* pathway, and in some comparative contexts covary with traits associated with sugar rewards. Yet the chemical ecological evidence remains uneven, and it is unclear whether these compounds function as evolved signals. A key question is therefore whether ripe fruits have evolved to signal ripeness and sugar level to mammal dispersers through synthesis of aliphatic esters.

The current study tests the hypothesis that fruits have evolved to utilize aliphatic esters as an honest signal to attract frugivorous seed dispersers. We addressed this hypothesis using a novel model system of 13 wild fig species (*Ficus*: Moraceae, Table S1) from Madagascar by integrating dispersal ecology derived from a novel interaction network with metabolomic analysis of fig scent, nutritional differentiation of sugars, and gene expression data obtained through RNA-seq. We conducted a comparative study on species dispersed primarily by lemurs, the local primates that are heavily olfaction-oriented and are likely to rely on chemical communication with fruits, and species dispersed by birds, a natural control group that is more visually oriented and relies less on olfactory cues, and is hence expected not to subject fruits to strong selection pressures to emit scent signals. We addressed the following predictions: (a) aliphatic esters are significantly more common in mammal-dispersed than in bird-dispersed *Ficus* species. Next (b), aliphatic-ester abundance is positively correlated with sugar concentration across multiple mammal-dispersed species, which are expected to benefit from signaling sugar levels. Finally, (c) plants actively generate ester signals by increasing *AAT* expression. Together, our results reveal that aliphatic ester production is associated with sugar content when mammal visitation occurs.

## Results

Across the full dataset, 25 distinct aliphatic esters were detected among the volatile samples from the 13 *Ficus* species (Table S2A,B,C). Two esters (ethyl acetate and methyl acetate) were detected in every species sampled in the dataset. While esters as a class are broadly present across *Ficus* fruit scent, individual ester identity and abundance differ substantially among species. Beyond the shared cores, aliphatic ester composition was highly species-specific with several compounds restricted to only one or two species and within species varied considerably.

### ***a.*** Mammal-dispersed fruits show higher aliphatic ester abundance

To test the prediction that aliphatic esters are more common in mammal-dispersed species, we used a linear mixed model using log transformations of aliphatic ester concentration and mammal visitation index, which is the proportion of mammals (primarily lemurs; see Materials and Methods on ecological networks) visiting fruits of each species (Figure S1). Across 13 *Ficus* species, we found a statistically significant positive association between the mammal visitation index and ester concentration (estimate = 3.41, SE = 0.67, df = 141, t = 5.13, p < 0.001, Figure 1). The model showed substantial individual-level variation, with 67.6% of the total variance explained by differences among individuals within species. Additionally, ester concentration showed marginally significant phylogenetic structuring across the species in our dataset (Pagel’s λ = 0.769, p = 0.038; Blomberg’s K = 0.564, p = 0.083, Figure S2).

**Figure 1.**
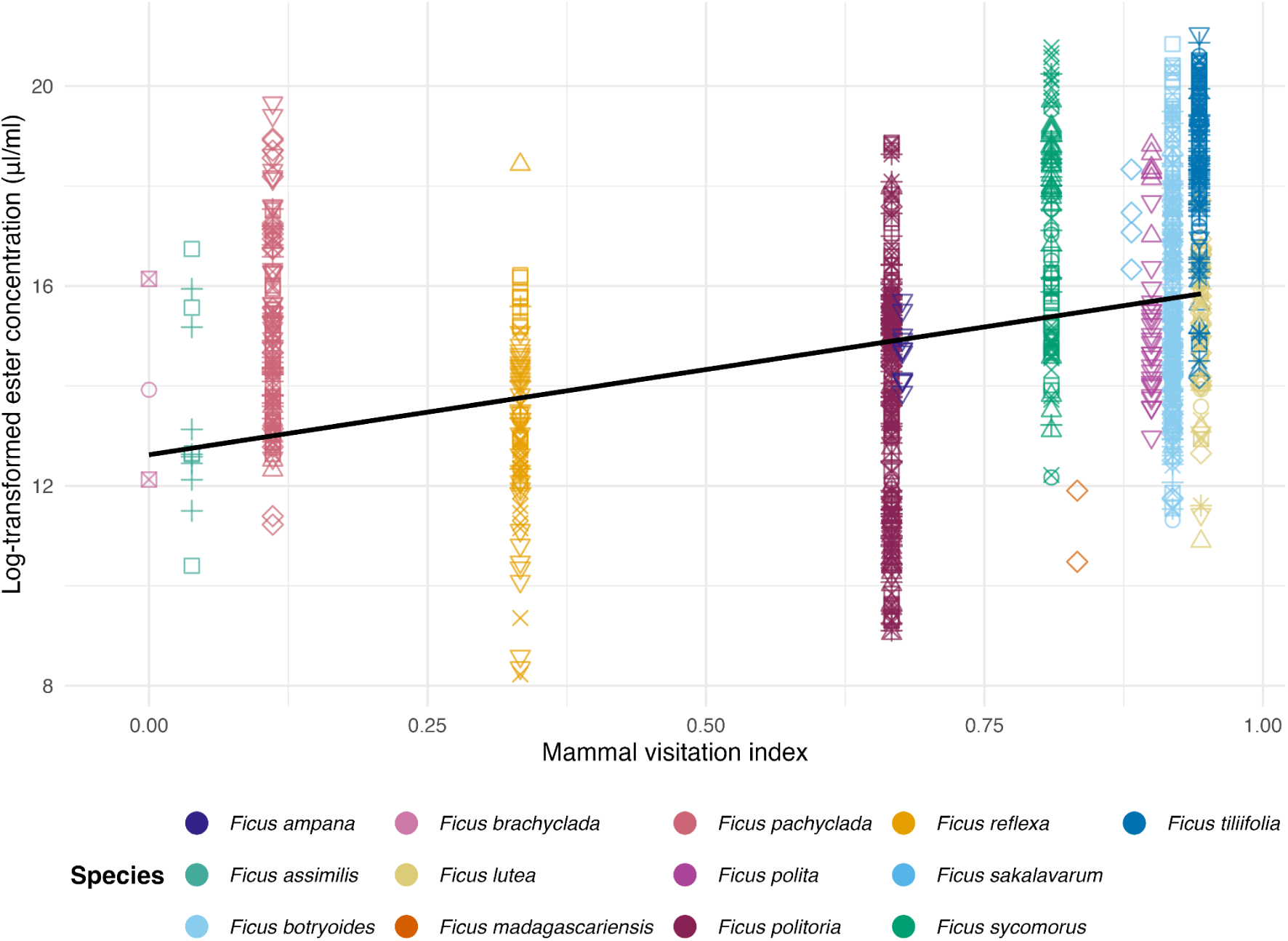
Species-level variation in aliphatic ester concentration explained by mammal visitation. Each dot represents one fruit sample, plotted by its mammal visitation index (x-axis) and log-transformed normalized ester concentrations (y-axis). Colors indicate the *Ficus* species of each sample, and unique symbols represent different individual trees. The black line shows the model-predicted fixed-effect relationship from a linear mixed model (estimate ± SE = 3.41±0.67, t141=5.13, p<0.001) with individuals nested within species as a random intercept; the intercept was set to the global average.

### ***b.*** Esters signify sugar levels, but only in mammal-dispersed species

We used a linear mixed-effects model (see Materials and Methods) to test whether ester concentration varied with sugar concentration and mammal visitation, that is, whether the correlation observed above was present only in species that benefited from attracting mammals. We found the degree to which esters and sugars are correlated depends on the main animal disperser (that is, the interaction term was statistically significant: slope = 1.69, SE = 0.77, p = 0.029, Figure 2). As mammal visitation increased, the positional relationship between sugar and ester increased (Figure 2). Sugar concentration showed no relationship with ester concentration in species mostly dispersed by birds.

**Figure 2.**
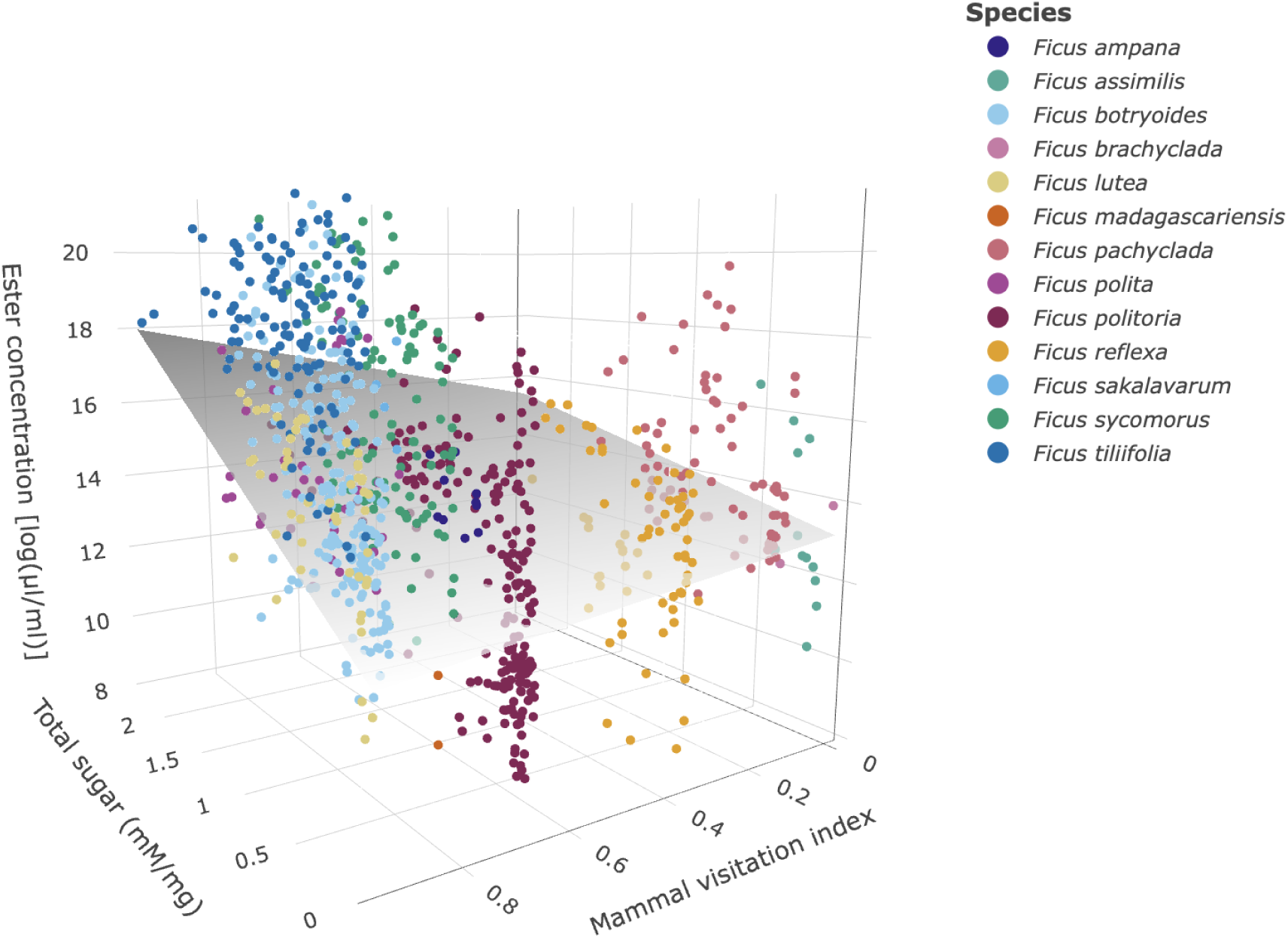
3D surface plot of the predicted ester concentration as a function of total sugar and mammal visitation index across 13 ripe fig species. Total sugar concentration (mM/mg), log-transformed ester concentration (log µl/ml), and mammal visitation index are plotted on the x-, y-, and z-axes, respectively. The surface represents model-predicted values from the mixed-effects model (fixed effects only), and colored points show observed values per species. The flat, horizontal ridge running along the surface when the mammal visitation index is 0 indicates that, in the absence of mammal visitors (bird-dominated visitors), ester concentration shows no meaningful relationship with sugar content. In contrast, as mammal visitation increased, the surface rose steeply, revealing that higher ester concentrations were associated with greater sugar content only in fruits that attracted mammal visitors. This interaction suggests that ester volatile production may function as an honest signal of reward quality, but only in a mammal visitation context.

### ***c.*** AAT activity is potentially correlated with sugar and ester levels

To further investigate whether plants actively produce aliphatic esters in the presence of sugars, we first identified candidate/marker genes responsible for ester synthesis. We used the genome-guided transcriptome of a single mammal-dispersed species (*Ficus tiliifolia*) to search for BAHD-family acyltransferases (Pfam: PF02458) and whose expression was associated with both sugar and esters. Among identified 67 BAHD family genes, “FICUtili007618t1” showed significant positive associations with both ester and sugar concentrations. To be specific, “FICUtili007618t1” was positively correlated with ester concentration (Spearman: ρ ≈ 0.46, *p_adj_* ≈ 0.013; Pearson: r ≈ 0.4, *p_adj_* ≈ 0.099) and also positively associated with sugar concentrations (Spearman: ρ ≈ 0.37, *p_adj_* ≈ 0.053; Pearson: r ≈ 0.41, *p_adj_* ≈ 0.028; Figures 3A and B, Table S2). Together, other two genes (“FICUtili007510t1,”and “FICUtili006443t1”) showed significant correlations. While “FICUtili007510t1” was moderately positively associated (Spearman: ρ ≈ 0.43, *p_adj_* ≈ 0.025 and Pearson: r ≈ 0.42, *p_adj_* ≈ 0.025, Table S2) with sugar concentration, “FICUtili006443t1” was negatively associated (Spearman: ρ ≈ –0.47, *p_adj_* ≈ 0.012 and Pearson: r ≈ –0.45, *p_adj_*≈ 0.023, Table S2) with sugar concentration (Figure 3A).

**Figure 3.**
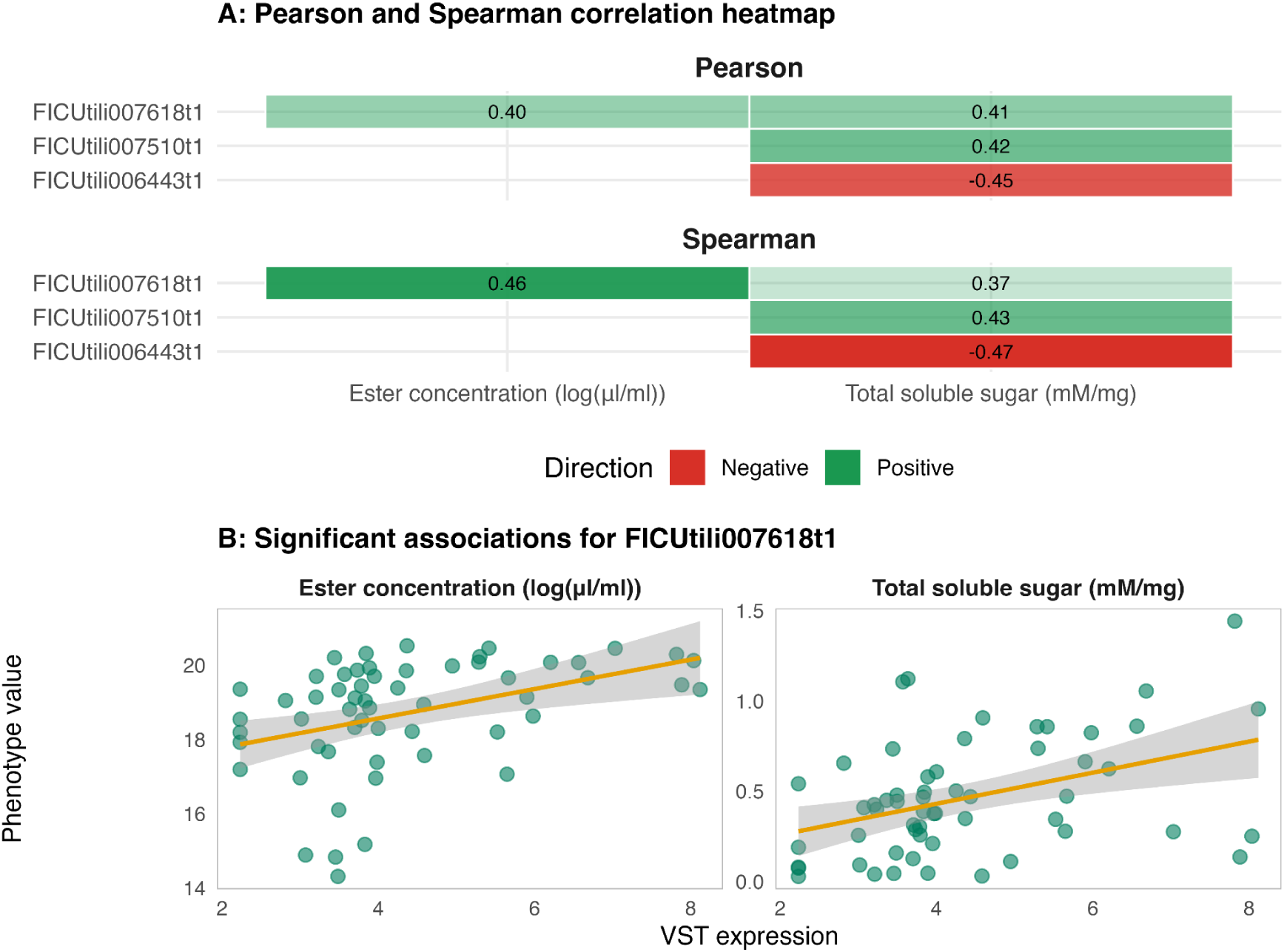
Statistically significant correlations between gene expression and *Ficus* phenotypes (A) and scatter plots of the relationship between variance-stabilizing transformation (VST), normalized gene expression and phenotype value (sugar content and ester concentration) for the gene “FICUtili007618t1” (B). (A) Tile color indicates the direction of the correlation (green for positive, red for negative) while color saturation reflects its strength, with more saturated tiles indicating stronger correlations and paler tiles indicating weaker ones. The corresponding correlation coefficient is printed within each cell. (B) Each point represents an individual sample, with gene expression plotted against the corresponding phenotype value. The line shows the fitted linear regression, and the shaded band represents the 95% confidence interval, illustrating the direction and strength of association.

Phylogenetic analysis placed “FICUtili007618t1” and “FICUtili007510t1” gene within a clade are known to utilize aliphatic alcohol-derived substrates, which are involved in/essential intermediate products of aliphatic ester biosynthesis. This indicates that “FICUtili007618t1” is likely to be the target *AAT* gene (Figure S3A,B). In addition, this clade was functionally characterized hydroxycinnamoytransferases including PtFHT1, AtASFT/HHT1, StFHT and AtFACT, which are involved in cutin and suberin metabolism. Several other *Ficus* genes (“FICUtili006946t1”, “FICUtili007200t1”, and “FICUtili007250t1”) clustered with experimentally characterized alcohol acyltransferases associated with volatile ester biosynthesis, including FvVAAT, RhAAT1 and CmAAT4 (Figure S3B). Together, these results identify a small suite of BAHD acyltransferases associated with fruit ester metabolism, although their precise biochemical functions remain to be determined.

## Discussion

Our objective was to test the hypothesis that attraction of olfaction-oriented mammalian seed dispersers, mostly lemurs, has driven fruits to actively synthesize esters as an honest signal of fruit sugar content. Our results reveal that (a) aliphatic esters are significantly more common in the fruit scent of ripe fruits of species that rely on dispersal by mammals, as opposed to those relying on visually oriented birds, and (b) aliphatic esters signify sugar levels in mammal-dispersed species, but not in bird-dispersed species. In addition, we identified (c) the candidate gene “FICUtili007618t1,” which likely encodes an *AAT* as it belongs to the BAHD acyltransferase and specifically clusters with amino acid sequence alignment known to accept aliphatic alcohols as a substrate, and its expression is positively correlated with sugar and esters in ripe fruits of a single model species (*F. tiliifolia*).

The aliphatic ester content increased with the increasing number of mammal visits. This indicates that esters are substantially more common in fruits that depend on olfaction-oriented mammals for dispersal. This increase in ester concentration in mammal-dispersed fruit species is consistent with previous studies that support greater ester abundance in mammal-dispersed fruits (17) and complements existing evidence on frugivores using chemical cues (17, 25). Mammal-dispersed (mammal-like) fruits exhibit higher ethanol concentrations than bird-dispersed fruits, implying that the metabolic pathways generating volatile esters may be upregulated in mammal-dispersed species (52). This strongly supports the hypothesis that, while esters are common in the fruits of many species, they are uniquely concentrated in fruit species that depend on chemical communication with seed dispersers. Given that ester concentrations show only marginal phylogenetic structuring across species, the dataset is consistent with broader patterns, indicating that fruit metabolites are shaped more by ecological and functional factors than by shared ancestry (53, 54). Mammal-associated variation between visitation and ester concentration remained robust after accounting for shared ancestry, consistent with the expectation that olfactory-guided dispersers impose direct selection on fruit chemical signals. Indeed, ecological and functional explanations are most likely, as selective pressures imposed by dispersers may drive differences in fruit traits, consistent with the dispersal syndrome hypothesis and disperser sensory ecology.

Notably, ester emissions showed large intraspecific variation (about 32.4%), especially among the more mammal-dispersed species, which had lower ester levels on average (Figure 1). Individual fruits with lower esters may be biochemically limited in their capacity to draw on precursors linked to sugars, as this relationship holds not only across species but also within individual trees (37). One possible interpretation is that fruit scent reflects differences in evolutionary strategy among plant species. Bird-dispersed plants may gain little from elaborate olfactory signaling and therefore invest relatively little in scent, whereas plants associated with mammal dispersers may invest to varying degrees, from modest to substantial scent production. Greater investment may also increase trade-offs with defense, resource allocation, and other functions of fruit chemistry. Ripe fruits, for example, retain secondary metabolites that can deter antagonists, inhibit pathogens or germination, alter gut retention, and otherwise influence dispersal outcomes; these functions can coexist but may constrain investment in attraction (55). Across fruit-dispersal systems, nutrient-rich fruits tend to be removed more readily, whereas phenolic defenses can reduce removal, illustrating this reward–defense trade-off (56). This interpretation is consistent with the observed interspecific pattern: bird-associated species showed relatively similar scent profiles, whereas lemur-associated species exhibited greater variation in both scent quantity and composition (17). Bird-associated fruits showed little change in scent production between unripe and ripe stages, whereas lemur-associated fruits showed larger increases; among lemur-associated species, greater increases in scent quantity were also associated with greater compositional shifts (16). Thus, when olfactory signaling provides little selective benefit, investment may remain low and relatively uniform across species. The resulting pattern is therefore not necessarily an anomaly: when a trait is weakly useful, it may be small and similar across species; when it becomes beneficial, variation can increase because species balance signaling benefits against costs, constraints, and competing chemical functions.

Esters were not only more common in mammal-dispersed species, but were also positively associated with sugar, particularly in those species. This indicates that where chemical communication is relevant, the ester fraction in fruit scent is an honest signal for fruit quality. Critically, the presence of sugar-ester link was only in mammal-dispersed species indicates that a *potential* link between sugars and esters is not inevitable. This implies that in mammal-dispersed species, where chemical communication is likely to offer fitness benefits to the plant, the fruit actively converts rewards (sugars) into signals (esters), thus generating an honest signal of fruit quality. Indeed, a necessary additional prediction to test here would be that frugivores use fruit scent to identify fruit quality. While not tested here directly, evidence that this is the case is available from lemurs in the same environment, feeding on figs and other fruit species (25). Across species, sugar availability consistently predicts ester production, suggesting that the link between ripening metabolism and scent signaling is extending beyond species-specific but reflects a conserved biochemical mechanism (33, 37). The link between fruit scent and mammalian dispersal is well established in this system (17, 25, 33, 37): mammals rely on scent to select fruit and are drawn to profiles indicating high sugar content. Our results add that the active conversion of reward (sugar) to esters (signal) is driven by plant-animal interactions. Scent therefore would function as a plant-controlled advertisement of reward, and its production should be maintained by selection through enhanced seed dispersal.

It is important to note that not all aliphatic esters originate from ethanol produced via sugar fermentation; a defined subset is the R-acid ethyl esters (e.g., ethyl R-ate) and the ethanoic-acid R-yl esters (e.g., R-yl acetate: ethyl acetate) (57–59). Nonetheless, in our dataset sugar fermentation esters are not just a majority subset, but they are also effectively the entire aliphatic ester signal (Table S2C). Their mean concentration accounts for 94.3% of the total mean aliphatic ester concentration, and at the individual sample level, this dominance is even more pronounced: the median sample exhibits 99.5% of its total aliphatic ester signal derived from sugar fermentation esters (Table S2C). As such, at least in this model system the ester signal includes practically only compounds whose synthesis is related to fruit quality. Due to non-specific AAT activity, an increase across ester classes is expected. The AAT activity model therefore predicts that more alcohol-derived esters should mean elevated enzyme activity, hence, more esters overall.

While these results indicate that the presence of esters in fruit scents results from the active conversion of sugar and its downproducts to the ester signal, the evidence they provide is indirect. The transcriptomic analysis provides a first glimpse into the molecular mechanisms underlying the association between sugar content and ester signalling. Presumably, increased ester production is allowed by the increased availability of alcohol and acyl donors resulting from higher sugar levels, leading to increased transcription of *AAT*s. Our transcriptomic analysis of a single species (*F. tiliifolia*) offers a potential mechanistic explanation through the correlations of contig “FICUtili007618t1” with sugars and esters with confirmation of its alignment with other *AATs*. Together, this is a strong indicator that it is the gene responsible for ester synthesis, including non-ethanol-derived esters, and that fruits with higher sugar levels *actively* increase its expression, yielding a higher ester signal. Previous evidence for rising sugar activities in *AAT* expression provides limited but suggestive clues. For example, *AAT* enzyme activity seems well correlated with total soluble solid content in Arava melons (60) and *AAT* gene expression was also detected throughout banana ripening (61). Other studies on some domestic fruits have indicated that the *AAT*-encoded enzyme controls C6-ester synthesis during ripening, providing a mechanistic basis for increased ester levels in ripe fruits or *AAT*-1 as a key regulator of ester formation in apples, illustrating how *AAT*s activity correlates with ester richness across species (62, 63).

Unexpectedly, the strongest expression-supported candidate, “FICUtili007618t1”, did not cluster with canonical fruit alcohol acyltransferases but instead grouped with BAHD hydroxycinnamoyltransferases involved in cutin and suberin biosynthesis (64–68). Members of this lineage contribute to the assembly of extracellular lipid polyesters that influence the physical properties of plant surfaces, including permeability and diffusion barriers. Their known activities attach to fatty alcohols, demonstrating an ability to utilize aliphatic alcohol-derived substrates as chemical currency to volatile forming. This phylogenetic placement opens an alternative mechanistic interpretation of the sugar-ester relationship. Volatile signalling depends not only on the synthesis of volatile compounds but also on their storage, retention and release from fruit tissues. The plant cuticle is increasingly recognized as an active regulator of volatile emission rather than a passive barrier, with cuticle composition and structure influencing the diffusion dynamics of hydrophobic volatile compounds (69, 70). Consequently, genes involved in cutin or suberin assembly may indirectly affect the strength of olfactory signals perceived by frugivores by modifying fruit surface layers and volatile permeability. An important caveat is that functional convergence is widespread among these genes: different species can independently recruit different genes to perform similar biochemical functions. This makes gene function difficult to infer from phylogenetic position alone. Indeed, AATs from different species are distributed broadly across the phylogeny of the gene family rather than forming a single, tightly clustered lineage. Thus, although our candidate genes may represent the same canonical AAT function, we cannot exclude this possibility because they derive from a non-model system in which gene function has not been experimentally characterized.

In this context, elevated expression of “FICUtili007618t1” in sugar-rich fruits could influence ester signalling in at least two non-mutually exclusive ways. First, the gene may participate directly in ester-associated metabolism through a specialized BAHD acyltransferase activity. Second, it may influence the retention or emission of volatile esters through effects on cuticle composition and permeability. Under either scenario, the positive association between “FICUtili007618t1” expression, sugar concentration and ester abundance indicates that ester signalling is actively regulated during ripening rather than arising passively from fruit chemistry. At the same time, phylogenetic analysis identified additional *Ficus* genes (“FICUtili006946t1”, “FICUtili007200t1” and “FICUtili007250t1”) that cluster with experimentally validated volatile alcohol acyltransferases such as FvVAAT, RhAAT1 and CmAAT4 (40, 71, 72). These genes therefore represent strong candidates for the direct biosynthesis of volatile esters, whereas “FICUtili007618t1” may represent a complementary mechanism influencing the deployment of the signal. Distinguishing between these possibilities will require biochemical characterization of substrate specificity, including both alcohol acceptors and acyl donors, as well as experimental assessment of effects on volatile production and emission.

In summary, by integrating data from dispersal ecology, metabolomics, and transcriptomics, our results strongly support the hypothesis that esters serve as an honest signal driving seed-dispersal interactions. They are common but are uniquely present in species that depend on chemical signals; they are an honest signal, but not universally so rather, only in the same species that rely on chemical communication. And there is indication that they are actively synthesized by plants rather than appearing spontaneously. The candidate gene has so far been identified in a single taxon, and its role in facilitating honest signaling of fruit quality remains unconfirmed. Both limitations point towards near future works on analyzing selection dynamics (e.g. dN/dS ratios) from candidate gene sequences across multiple species, testing the prediction that dispersal by mammals generates positive or stabilizing selection, whereas dispersal by birds may allow loss of function. These current results demonstrate that while chemical communication is a medium through which plant-frugivore interactions are mediated, aliphatic esters are the language and content of that communication, driving critical seed-dispersal interactions and hence plant reproduction, forest regeneration, and ecosystem functioning.

## Materials and Methods

### Fieldwork and sample collection

#### Ecological network

We monitored 52 individual *Ficus* trees across 4 sites using 10 camera traps (mounted at a mean height of 4.9 ± 4.2 m, Browning Spec Ops Elite HP4 Trail Camera, Birmingham, Alabama, United States) redeployed sequentially as fruits ripened, yielding a mean deployment duration of 36.5 ± 22.5 days (range: 2–83 days) per tree between April 2022 and November 2024. From these deployments, we derived a mammal visitation index for each species, the proportion of independent feeding events attributable to mammals (lemurs) versus birds, which served as our primary proxy for frugivory intensity, guild composition. Visitation records were classified into two guilds, mammal and bird. The mammal guild was dominated by lemur sightings (92.7% of mammal records) but also included "small mammal" (6.3%), carnivora (0.8%), and unspecified "mammal" (0.2%). Otherwise, butterfly, reptile, or unidentified guild were excluded from the visitation index due to low sample size or ambiguous identification. The mammal visitation index was calculated by the number of mammal visits per total visits (mammal plus bird visits) for each species. The range of value was from 0 to 1, in which 0 indicates zero percent of mammal visits (all bird visits) and 1 represents 100% mammal visits (no bird visits) (Figure S1).

### Sample collection

Fieldwork was conducted in 2022 in four sites representing Northern, Eastern, Western, and Southern Malagasy forest: (a) Talatakely region, Ranomafana National Park (RNP) managed by Madagascar National Parks, (b) Andasibe-Analamazaotra Special Reserve (AD) managed by Mitsinjo, (c) Ankarafantsika National Park (AK) managed by Madagascar National Park and (d) Kirindy Forest (KIR) managed by CNFEREF. *Ficus* species are among the most diverse and ecologically important plants in tropical forests, where figs act as keystone food resources for a wide range of vertebrates (73). We collected ripe fig samples from 13 species of *Ficus* (Moraceae) opportunistically across Madagascar (Table S1). Ripeness was determined based on specific external color, fruit softening, and the presence of mature seeds, confirmed by opening fruits after sampling. Fruits were collected using disposable, nitrile gloves (Th. Geyer GmbH, Renningen, Germany), only in the morning local time, directly from trees, and transported to the laboratory within 3 hours. Fruits (1 – 10 fruits) from the same individual were pooled and normalized to be treated as a single biological replicate. For each tree, individual fruits were paired for simultaneous scent sampling and sugar analysis, with the remaining fruits allocated to RNA extraction.

### Fruit scent analysis

Fruit scent was collected using a semi-static headspace sampling protocol (17, 33). Ripe fruits were enclosed in heat-resistant oven bags (40 cm; Toppits, Minden, Germany). One end of the bag was sealed with a zip tie, and the other end was tightened around a Teflon tube connected to a volatile trap probe that consisted of quartz tubes containing Tenax TA, Carbotrap, and Carbosieve III (Sigma-Aldrich, Taufkirchen, Germany), held in place by glass wool. Fruits (Table S1) were in the bags for 20 min, after which headspace air was drawn through the trap for 15 min at a flow rate of 200 ml/min using a pump. After sampling, traps were sealed in Teflon-capped glass vials, wrapped with Parafilm, brought back to Germany until analysis, and stored at -20 °C. Negative control samples were collected daily after all samples using empty bags processed identically.

VOC samples were analyzed using a Gas Chromatography of Shimadzu Nexis GC-2030, with Mass spectrometry called Shimadzu GCMS-QP2020 NX system (Shimadzu, Canby, Oregon, USA) equipped with a TD30-R thermal desorption unit (Shimadzu, Nakagyo-ku, Kyoto, Japan) and an Shimadzu DB5 capillary (SH-I-5MS: 5 % diphenyl/ 95 % dimethyl polysiloxance with low polarity) column (30 m x 0.25 mm x 0.25 µm). Thermal desorption was performed by heating traps with tube desorption at 300 °C, 60 ml/ min for 15 min. Trap cooling temp -20 °C, trap desorb 250 °C, joint at 200 °C, and both valve and transfer line at 230 °C. Subsequently injected in splitless and split mode with ratio 1:1, 1:40, 1:50, and 1:70 due to the high intensity of VOCs. The oven temperature program started at 45 °C (hold 3 min), increased at 10 °C per min to 325 °C and held for 20 min. The MS operated in electron ionization mode (35-450 m/z scan from 1.10 min to 48.5 min). Transfer line and ion source temperatures were set to 280 °C and 230 °C, respectively.

VOCs were identified with the Shimadzu LabSolutions software and retention indices calculated using an n-alkane standard (C8-C20). Compounds detected in negative controls, in system controls (empty stainless steel tubes), were considered contaminants and excluded. External standard mixtures were run separately and regularly during running the scent sample with identical settings and split modes. The calibration curves include 2 µl/ml of methyl hexanoate (≥99%), methyl butyrate (≥98%), ethyl butyrate (≥98%), methyl acetate (≥98%), ethyl hexanoate (>98%), and ethyl acetate (≥99%) all in Diethyl phthalate (99.5%) from Sigma-Aldrich (Taufkirchen, Germany). This approach assumes comparable calibration slopes, detector response factors among compounds eluting within similar retention time windows that have similar chemical structure. Calibration curves were generated using multiple concentration levels of standards across the analytical range and fitted by linear regression of peak area versus concentration corresponding to the matching split ratio. For compounds lacking direct calibration standards, concentrations were estimated using the calibration curve of a representative standard with a similar retention time (e.g., butanoic acid ethyl ester equivalents for compounds eluting between 2.868 and 4.878 min). All VOCs in a sample were quantified by converting the absolute intensity amount to concentrations (µl/ml) through external calibration standard curves and then normalizing these concentrations by the number of fruits in the samples (17). The functional unit in this study is the fruit truss, as scent was measured from the entire truss because the animal will perceive and decide based on the scent emitted by the entire fruit truss. However, trusses varied in the number of individual fruits, scent measurements were normalized by the number of fruits per truss to account for differences in fruit number.

### Sugar analysis

After scent collection, part of the fruit tissue was oven-dried at 45-50 °C for at least 24 h and stored with silica gel during transportation to Germany until analysis (74). Dried samples were ground to a fine powder using a tissue homogenizer (Retsch MM400, Haan, Germany). Sugar extraction followed a modified protocol from Sławińska et al. (2021). Briefly using 100 mg of dried powder was extracted with 400 µL of 80% ethanol (Merck, Germany), incubated at 80 °C for 30 to 40 mins and centrifuged at 5000 x g for 15 min (Heraeus Multifuge X1R Centrifuge, ThermoFisher Scientific GmbH, Osterode am Harz, Germany). Supernatants were diluted to 1 mL with 80% ethanol. A 100 µL aliquot was added with 300 µL acetonitrile (ACN, for HPLC LC-MS grade, VWR chemicals, France) and incubated at -20 °C overnight, followed by centrifugation at 16,000 x g for 15 min. The supernatant was transferred to Liquid Chromatography vials.

Sugars were separated on a Dionex Ultimate 3000 UHPLC system using a precolumn Shodex Asahipak NH2P-50G 4A (5 µm, 4.6 × 10 mm) connected to a Shodex Asahipak NH2P-50 4E (5 µm, 4.6 × 250 mm, Japan). The compounds were detected with a Corona Veo RS charged aerosol detector (Germering, Germany) and a photodiode Array Detector (PDA; Ultimate 3000 series system DAD-3000 (RS), Thermo Fisher Scientific, Waltham, MA, USA). The mobile phase consisted of 75% ACN to 25% water at a flow rate of 1ml/min. The injection volume was 10 µL, and the total run time was 22 min.

Data analysis was performed using Chromeleon 7 Data System software (Thermo Fisher Scientific, version 7.2 SR4, Waltham, MA, USA). Individual sugars were qualified using external calibration curves with standards including D-(-)-fructose, D-(+)-glucose, and D-(+)-saccharose (sucrose) from Carl Roth GmbH (Karlsruhe, Germany). A 2 mg/mL stock solution was diluted with water to prepare working standards at different concentrations. Peak intensities detected in blanks (ACN only) and negative controls (extraction without sample material) were excluded as contaminants. The identified sugars contained fructose, glucose, and sucrose, while other unidentified or uncertain annotations were grouped as “others” (arabinose, xylose, galactose, mannose, and their corresponding sugar alcohols, as they shared similar retention time) but were categorized as carbohydrates based on their UV absorbance signals. The total sugar concentration used in this study was the sum of three main sugars (fructose, glucose, and sucrose).

### Transcriptomics

After scent and sugar collection, the remaining part of the fruit tissue was submerged in liquid nitrogen and kept at -80 °C until transportation to Germany. We selected *Ficus tiliifolia* as our model species because it has been characterized as a lemur specialist and has been shown to contain aliphatic esters in its ripe fruits, as well as to exhibit a positive correlation with total sugar content (17, 37). 57 samples of *Ficus tiliifolia* were stored in a cryo vapor shipper with a hard shell (CX100, Worthington Industries, Columbus, Ohio, USA), then transported by air to Germany.

Frozen tissue was ground to a fine powder in liquid nitrogen using a mortar and a pestle. Total RNA was extracted using a modified method in a two-day extraction from Reid et al. (2006). Sample powder with 300 mg was added to 6 mL of pre-warmed (65 °C) extraction buffer made by 300 mM Tris(hydroxymethyl)aminomethane hydrochloride, 25 mM Ethylenediaminetetraacetic acid, 2 M Sodium Chloride, 2 % Cetyltrimethylammonium bromide, 2 % Polyvinylpolypyrrolidone, 0.05% spermidine trihydrochloride, and just before using the extraction, adding 2 % beta-mercaptoethanol. Mixtures were vortexed, then incubated in a water bath at 65 °C for 10 min with occasional shaking. Fruit Mate (600 µL) was added, mixed by inversion, and samples were centrifuged (5,500 × g, 10 min, 4°C, Heraeus Multifuge X1R Centrifuge, ThermoFisher Scientific GmbH, Osterode am Harz, Germany). The supernatant was transferred to a new tube and extracted with 6mL chloroform: isoamyl alcohol (24:1), followed by centrifugation (5,500 × g, 15 min, 4°C). The aqueous phase was transferred and clarified by an additional centrifugation step (5,500 × g, 20 min, 4°C). Nucleic acids were precipitated by adding 600 µL 3 M NaOAc and 3.6 mL isopropanol, vortexing, and incubating at −80 °C for 30 min, followed by centrifugation (5,500 × g, 30 min, 4°C). Pellets were dissolved in 1–2 mL TE buffer, transferred to fresh tubes, and RNA was selectively precipitated by adding 300 µL 8 M LiCl and incubating overnight at 4°C. RNA was pelleted by centrifugation (5,000 × g for 5 mL tubes, 30 min, 4°C), washed twice with ice-cold 70% ethanol, air-dried, and resuspended in 70–100 µL Milli-Q water. Samples were stored at −80 °C. Assessment of RNA quality and quantity was done by 1 % agarose gel and NanoPhotometer P330 (Implen, Munich, Germany).

Library preparation and sequencing of the samples were performed by Azenta/GENEWIZ (Leipzig, Germany) using poly(A)-enriched libraries multiplexed and loaded onto the flow cell of an Illumina NovaSeq 6000/Xplus instrument according to the manufacturer’s instructions. The samples were sequenced using a 2x150 Pair-End (PE) configuration v1.5. Image analysis and base calling were conducted by the NovaSeq Control Software v1.7 on the NovaSeq instrument. Raw sequence data (.bcl files) generated from Illumina NovaSeq was converted into *fastq* files and de-multiplexed using Illumina bcl2fastq program version 2.20. One mismatch was allowed for index sequence identification. Samples received about 20 million paired-end reads. Raw data FASTQ went through a series of workflows under the High-Performance Computing (HPC) cluster at the German Centre for Integrative Biodiversity Research (iDiv). Raw sequencing reads were quality trimmed using *fastp* (76). Genome-guided alignment was performed with *HISAT2* (77) using the *Ficus hispida* reference genome (PRJCA016767; 68.84% read mapping). Transcriptome assemblies were generated with *Trinity* (78), yielding a BUSCO completeness score of 90.8%. Primary gene models were extracted from the aligned assemblies using *EviGene* (79), with subsequent BUSCO assessment indicating 93.5% completeness. Assemblies were functionally annotated using *MapMan4* (80), and transcript abundance was quantified with *Bowtie2* (81) and *Subread* (82).

For targeted analysis of the BAHD acyltransferase gene family, amino acid sequences from the *EviGene* output were screened using *HMMER*-*hmmsearch* (83) to identify proteins containing the PF02458 domain. BAHD-related sequences were extracted from the assembled transcriptome, combined with reference BAHD sequences (46), and used for phylogenetic reconstruction with *IQ-TREE* version 2.3.5 (84).

### Statistical analyses

All the analysis was carried out in R version 4.5.1 (85) using packages for phylogenetic comparative methods (ape (86), phytools (87)), mixed-effects modelling (lme4 (88), lmerTest (89), brms (90)), differential expression analysis (DESeq2 (91), edgeR (92), limma (93)), and visualization (ggplot2 (94), plotly (95)). Three predictions were tested: (a) aliphatic esters were more commonly found in mammal-dispersed species, (b) aliphatic esters were positively significant to the sugars across multiple species, and (c) candidate genes that predict *AAT* genes in mammal-dispersed fruits whose expression correlated with ester concentration and sugar concentration.

To test whether aliphatic esters were common in mammal-dispersed species (a), we used a mixed effect model, *lmer* (88), with ester concentration (log transformed) and mammal visitation with a random intercept effect structure of individuals nested in species. Next (b), we used a linear mixed-effects model with an interaction term, fitted using *lmer* (88) to test whether ester concentration as a response variable varied with sugar concentration and mammal visitation across multiple species. Fixed effects included total sugar concentration (sum of three sugars), mammal visitation index, and their interaction, with the same nested random effects structure as before (individual within species). To account for non-independence, we included random intercepts for individuals nested within species. The dataset included 873 observations from 133 individuals across 13 species.

Lastly, to pinpoint the candidate markers (c) predicting the *AAT* gene (in the BAHD acyltransferase family gene) in mammal-dispersed fruit, gene expression was normalized with variance stabilizing transformation (VST) from *DESeq2* (91). Before applying VST normalization, low-count genes were filtered to reduce noise using a threshold of at least 5 reads in at least 2 samples. Then, BADH gene extraction was performed to subset to only ester synthesis candidates. Correlation analysis was used to test which BAHD genes correlate with ester concentration (log-transformed) and total sugar concentration in both Pearson (linear) and Spearman (rank-based) analyses, with multiple testing correction using Benjamini-Hochberg FDR applied. Phylogenetic signal in mean ester concentration was estimated across species using Pagel’s lambda and Blomberg’s K (phylosig, phytools), with branch lengths computed. Both statistics were tested against a null of no phylogenetic structure (K: 999 permutations).

## Supporting information

Supplemental_all

## Acknowledgments

The project was carried out under research permits (Autorisation de recherche) for two field seasons of N° 383/21/MEDD/SG/DGGE/DAPRNE/SCBE.Re and N° 253/22/MEDD/SG/DGGE/DAPRNE/SCBE.Re. Export permits for samples were N°035N-EV02/MG21 and N°327N-EV12/MG22. We gratefully acknowledge the support of Andasibe–Analamazaotra Special Reserve (managed by Mitsinjo), Ankarafantsika National Park (managed by Madagascar National Parks), Kirindy Forest (managed by CNFEREF), and Ranomafana National Park (managed by Madagascar National Parks), as well as the staff of all parks, Centre ValBio, MICET, and the Mad Dog Initiative, for their invaluable assistance during our field seasons. The study was supported by a DFG (German Science Foundation) grant NE 2156/3-1. We also gratefully acknowledge the support of the German Centre for Integrative Biodiversity Research (iDiv) Halle-Jena-Leipzig, funded by the DFG (FZT 118, 202548816). Further, we thank Dr. Alexander Weinhold and Dr. Andreas Schedl for their support with analytical chemistry, and Dr. Renske Onstein for her advice on phylogenetic work.

