## Supplemental_all for "Fruit scent chemistry: adaptation to seed dispersal interactions"

**Figures and Tables Supplementary**

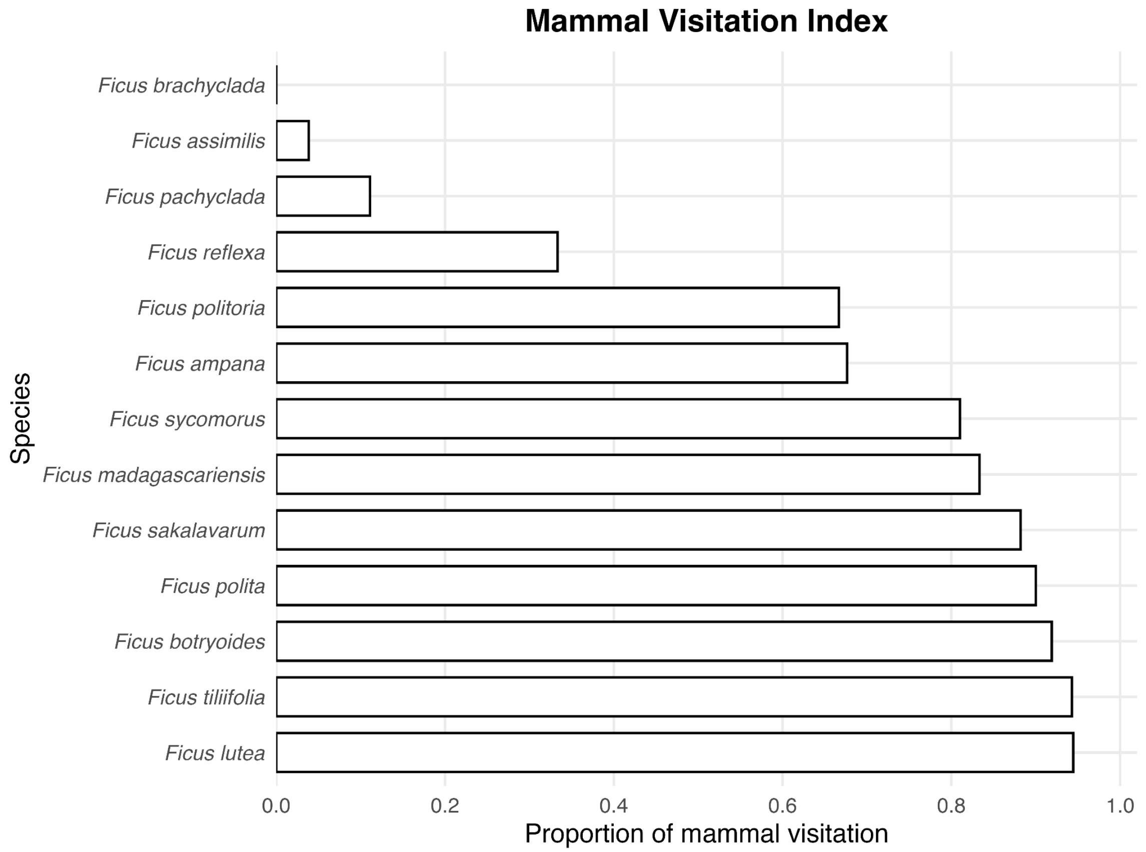


**Figure S1**. Mammal Visitation Index for different Malagasy *Ficus* species. Each bar represents one species, and the length illustrates the proportion of mammal visitation in comparison to total recorded visitation. The bar is in order by the lowest to the highest value of the mammal visitation index.

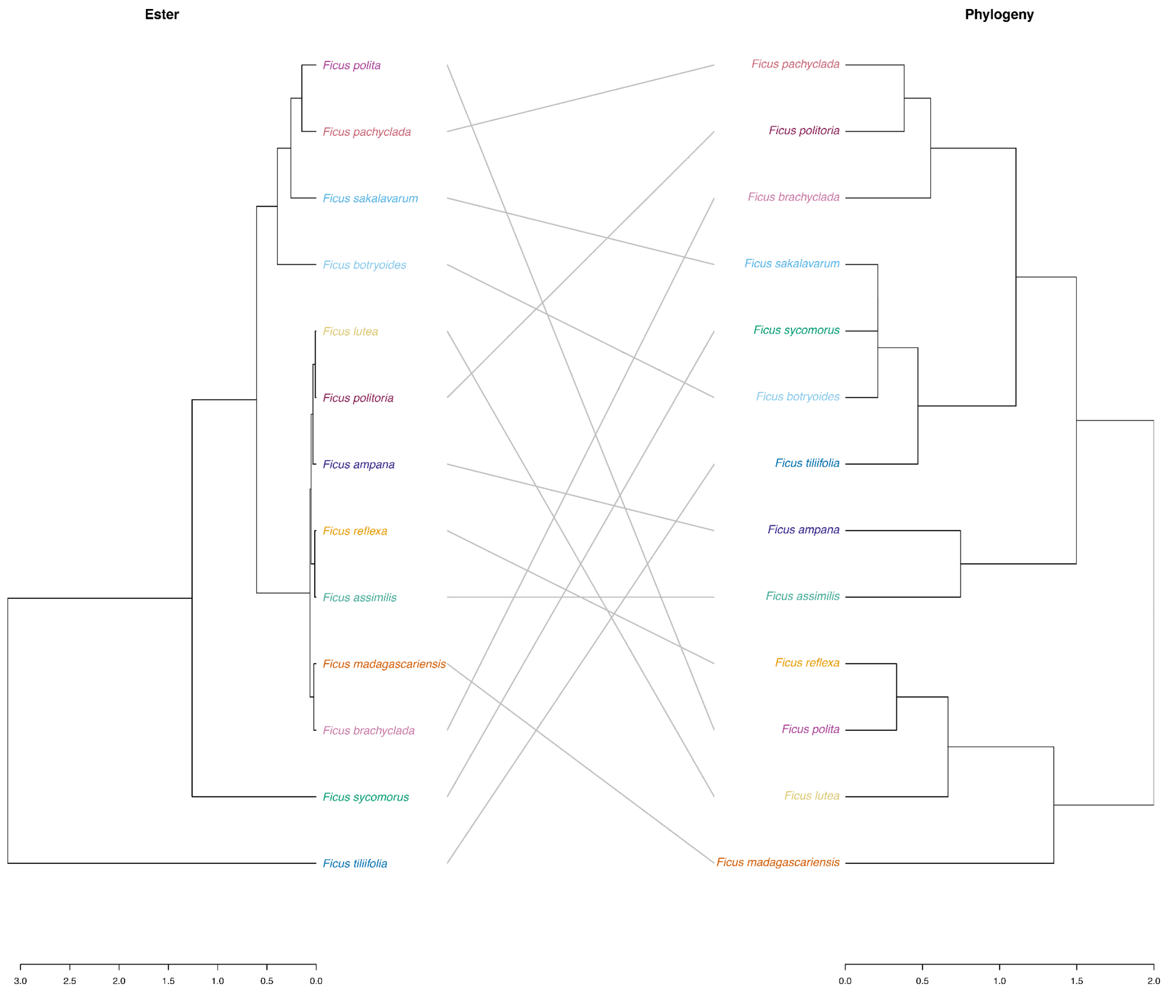


**Figure S2.** Tanglegram comparing a complete-linkage dendrogram of species-averaged fruit aliphatic ester profiles with the *Ficus*phylogenetic tree (phylogenetic signal: Pagel's $\lambda = 0.769$, $p = 0.038$; Blomberg's $K = 0.564$, $p = 0.083$).

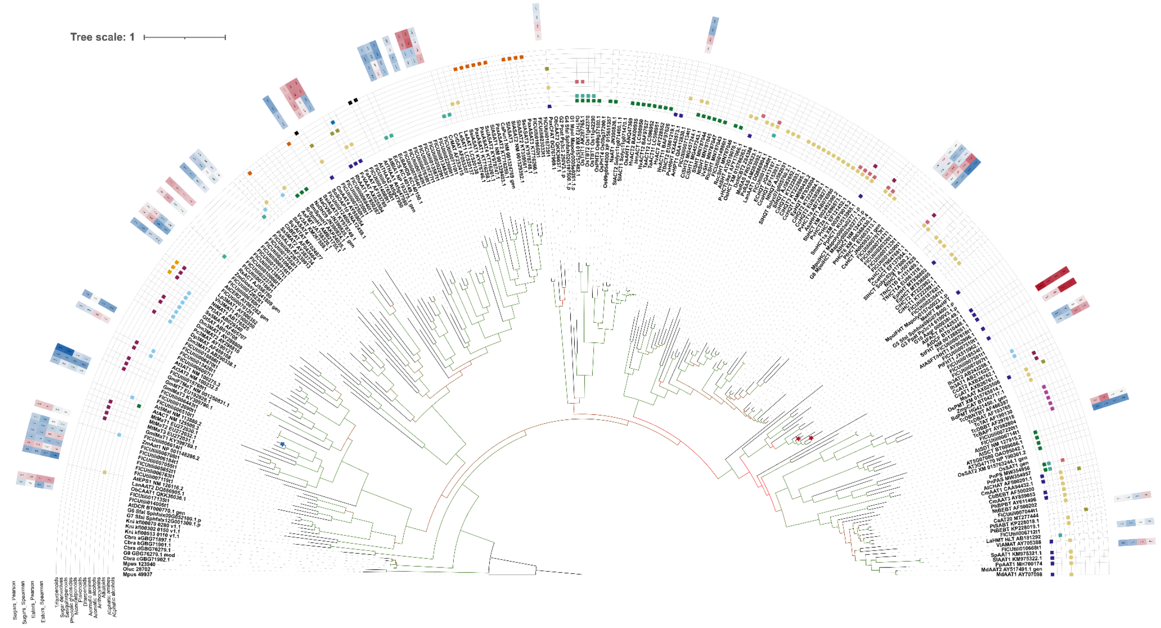


**Figure S3A.** Half-circle plot including annotations of substrate classes and correlations between BADH genes and phenotypes (esters and sugars). Stars in the branch indicate statistically significant tests. Blue indicates negative correlations, while red indicates positive correlations. Branch lengths are colored according to bootstrap support.

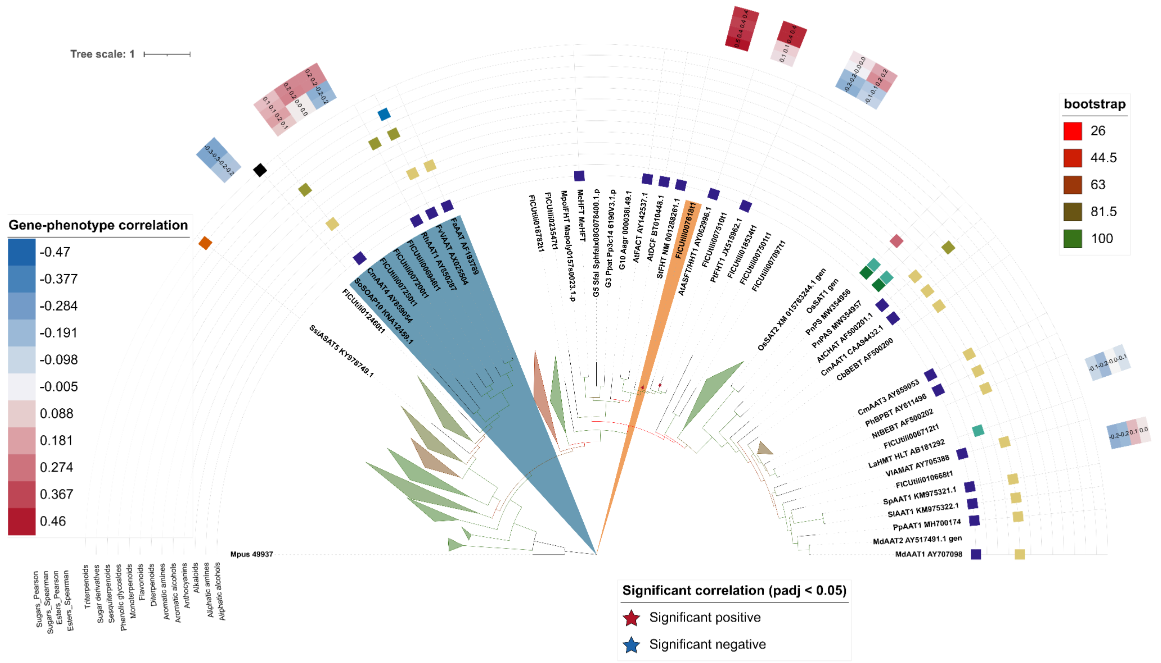

**Figure S3B.** The zoomed-in view of the half-circle plot with node collapse shows the focused branches containing "FICUtili007618t1" (orange), which exhibits the strongest association with ester and sugar concentrations in the expression analyses. The blue-highlighted clade contains "FICUtili006946t1", "FICUtili007200t1", and "FICUtili007250t1", which cluster with functionally characterized alcohol acyltransferases such as "FvVAAT", "RhAAT1", and "CmAAT4". The two branches distinguish the strongest expression-based candidate (orange) from the strongest phylogenetically supported candidates (blue).

**Table S1.** Summary of sample size in species, number of individuals, samples and mean samples per individual with standard deviation across 4 different field site Andasibe-Analamazaotra Special Reserve (AD), Ankarafantsika National Park (AK), Kirindy Forest (KIR), and Ranomafana National Park (RNP)

| **Species** | **Field site** | **Number of individuals** | **Number of samples** | **Sample per individual (mean ± SD)** |
|---|---|---|---|---|
| ***Ficus ampana* C.C. Berg** | AD | 1 | 11 | 11 |
| ***Ficus assimilis* J. Linn** | AK | 3 | 13 | 4.3 ± 3.5 |
| ***Ficus botryoides* Baker** | AD, AK, RNP | 33 | 262 | 7.9 ± 6.5 |
| ***Ficus brachyclada* Baker** | AD | 2 | 3 | 1.5 ± 0.7 |
| ***Ficus lutea* Vahl** | AD, RNP | 10 | 63 | 6.3 ± 7.4 |
| ***Ficus madagascariensis* C.C. Berg** | AK | 1 | 2 | 2 |
| ***Ficus pachyclada* Baker** | AK, RNP | 11 | 111 | 10.1 ± 7.5 |
| ***Ficus polita* Vahl** | AK, RNP | 2 | 33 | 16.5 ± 13.4 |
| ***Ficus politoria* Lam.** | AD, RNP | 43 | 246 | 5.7 ± 4.8 |
| ***Ficus reflexa* Thunb.** | AD, RNP | 11 | 84 | 7.6 ± 7 |
| ***Ficus sakalavarum* Baker** | AK | 1 | 4 | 4 |
| ***Ficus sycomorus* L.** | AK, KIR | 11 | 111 | 10.1 ± 7.8 |
| ***Ficus tiliifolia* Baker** | AD, RNP | 17 | 131 | 7.7 ± 7.1 |

**Table S2.** Volatile scent composition and ester dominance across *Ficus* species

A. Overall contribution of esters versus other volatile compound classes to the total scent blend, summed across all species and samples

| | Total concentration (µl/ml) | % in the total concentration |
|---|---|---|
| Aliphatic esters | 1329.41 | 54.31 |
| Others | 1118.40 | 45.69 |

B. Volatile compound richness and ester dominance per *Ficus* species, showing the total number of compounds detected, the number of aliphatic esters detected, and the percentage of the total volatile blend composed of esters

| Species | Number of detected compounds | Number of Aliphatic esters detected | Ester concentration in total scent (%) |
|---|---|---|---|
| *Ficus ampana* | 60 | 11 | 13.38 |
| *Ficus assimilis* | 49 | 7 | 27.10 |
| *Ficus botryoides* | 178 | 25 | 41.59 |
| *Ficus brachyclada* | 39 | 5 | 4.57 |
| *Ficus lutea* | 130 | 17 | 20.22 |
| *Ficus madagascariensis* | 23 | 3 | 0.93 |
| *Ficus pachyclada* | 158 | 21 | 25.08 |
| *Ficus polita* | 101 | 16 | 28.88 |
| *Ficus politoria* | 169 | 20 | 17.56 |
| *Ficus polyphlebia* | 108 | 21 | 40.61 |
| *Ficus reflexa* | 147 | 19 | 22.89 |
| *Ficus sakalavarum* | 49 | 8 | 27.79 |
| *Ficus sycomorus* | 155 | 26 | 64.33 |
| *Ficus tiliifolia* | 143 | 24 | 75.52 |

C. A list of aliphatic esters presented, showing the number of *Ficus* species in which each ester was detected and its mean concentration when present

| Aliphatic ester | Number of species detected | Mean ester concentration (µl/ml) |
|---|---|---|

| | | |
|---|---|---|
| comp_627_Acetic acid ethyl ester | 14 | 0.6814 |
| comp_661_Acetic acid methyl ester | 14 | 0.1490 |
| comp_320_Butanoic acid ethyl ester | 12 | 1.5413 |
| comp_693_2-Nonanol, acetate | 11 | 0.1126 |
| comp_250_Benzoic acid, 4-ethoxy-, ethyl ester | 11 | 0.0263 |
| comp_838_Hexanoic acid ethyl ester | 11 | 0.0186 |
| comp_637_Butanoic acid methyl ester | 10 | 0.0917 |
| comp_550_Acetic acid, butyl ester | 10 | 0.0779 |
| comp_73_Butanoic acid, 1-methylhexyl ester | 10 | 0.0186 |
| comp_902_Hexanoic acid methyl ester | 9 | 0.0875 |
| comp_169_Octanoic acid, methyl ester | 9 | 0.0383 |
| comp_102_2-Pentanol, acetate | 8 | 0.0249 |
| comp_850_Octanoic acid, ethyl ester | 8 | 0.0182 |
| comp_6_Heptadecanoic acid, ethyl ester | 8 | 0.0140 |
| comp_238_Nonanoic acid, methyl ester | 8 | 0.0077 |
| comp_232_Dodecanoic acid, 1-methylethyl ester | 8 | 0.0051 |
| comp_281_2-Butenoic acid, methyl ester, (E)- | 7 | 0.0438 |
| comp_528_2-Butenoic acid, ethyl ester | 7 | 0.0228 |
| comp_801_n-Valeric acid cis-3-hexenyl ester | 7 | 0.0125 |
| comp_222_1-Methylbutyl butyrate | 6 | 0.0708 |
| comp_24_Acetic acid, hexyl ester | 6 | 0.0597 |
| comp_211_2-Butenoic acid, ethyl ester, (Z)- | 6 | 0.0389 |
| comp_543_2-Hexenoic acid, ethyl ester | 4 | 0.0652 |
| comp_45_Oxalic acid, propyl undecyl ester | 3 | 0.0343 |
| comp_287_Heptadecyl acetate | 2 | 0.0060 |

**Table S3**. Statistically significant Pearson and Spearman correlation between GeneID and different phenotypes.

| GeneID | phenotype | rho Spear | P Spear | rho Pear | P Pear | Padj Spear | Padj Pear |
|---|---|---|---|---|---|---|---|
| **FICUtili006443t1** | Total sugar | -0.47 | 0.00024 | -0.45 | 0.00047 | 0.012 | 0.023 |
| **FICUtili007510t1** | Total sugar | 0.43 | 0.00102 | 0.42 | 0.00102 | 0.025 | 0.025 |
| **FICUtili007618t1** | Total sugar | 0.37 | 0.00429 | 0.41 | 0.00172 | 0.053 | 0.028 |
| **FICUtili007618t1** | Ester conc (log) | 0.46 | 0.00027 | 0.40 | 0.00201 | 0.013 | 0.099 |
